# Hepatocyte Growth Factor and β1-integrin signalling axis drives tunneling nanotube formation in A549 lung adenocarcinoma cells

**DOI:** 10.1101/2022.12.01.517334

**Authors:** Griselda Awanis, Sathuwarman Raveenthiraraj, Robert Johnson, Jelena Gavrilovic, Derek Warren, Anastasia Sobolewski

## Abstract

Tunneling nanotubes (TNTs) are thin cytoplasmic protrusions involved in long-distance cellular communication. The presence of TNTs has been found *in vivo* and *in vitro* studies in non-small cell lung cancer (NSCLC). Cancer cells transport a range of organelles and signalling molecules along TNTs, to confer a survival phenotype for the recipient cell, contributing toward chemoresistance and malignancy. Despite its important role in cancer progression, the molecular mechanisms underlying TNT formation is not well defined. Within the tumour microenvironment (TME) of NSCLC, hepatocyte growth factor (HGF) and its receptor, c-Met, are mutationally upregulated causing growth, and invasion. In this study, we report a novel crosstalk between HGF/c-Met and β1-integrin involved in the formation of functional TNTs in A549 cells. Through pharmacological inhibitor studies, we discovered Arp2/3 complex, MAPK and PI3K pathways were activated downstream of this crosstalk signalling axis. Furthermore, paxillin was recruited during this key process, localising at the protrusion site of HGF-induced TNTs, and therefore serving as the central link between the upstream and downstream regulators involved. Overall, these results demonstrate a novel strategy to inhibit TNT formation in NSCLC through targeting the HGF/c-Met and β1-integrin signalling axis, thus highlighting the importance of personalised multi-drug targeting in NSCLC.

## Introduction

Non-small cell lung cancer (NSCLC) accounts for 85% of all lung cancer cases. It is associated with high mortality rates as the majority of patients are diagnosed with advanced metastatic cancer (Molina et al. 2008). A vital aspect underlying growth and metastasis in cancer is the cellular communication existing between cells in the tumour microenvironment. Tunneling nanotubes (TNTs) are novel structures implicated in long-range cell-to-cell communication. TNTs are non-adherent, open-ended F-actin-based cytoplasmic protrusions and were initially discovered in cultured rat pheochromocytoma PC12 cells (Rustom et al. 2004). TNTs are characteristically long and thin structures, spanning up to 131μm in length and between 0.4-1.5μm in width in lung cancer cells (Wang et al. 2021). They display variability in their cytoskeletal composition, with microtubules present in the thicker TNTs (Önfelt et al. 2006). TNTs have been observed in multiple cell lines (Ariazi et al. 2017), including A549 lung adenocarcinoma cells (Wang et al. 2021; Dubois et al. 2018; Kumar et al. 2017; Wang et al. 2012). *In vivo*, TNTs were observed in human lung adenocarcinoma and mesothelioma tumour samples, thus highlighting their presence in solid tumours (Lou et al. 2012). TNTs facilitate the transport of cellular organelles such as mitochondria, Golgi vesicles, miRNA and signalling molecules to distant non-adjacent cells (Wang and Gerdes 2015; Wang et al. 2011; Lou et al. 2012; Lu et al. 2019; Thayanithy et al. 2014), consequentially promoting cancer progression and propagating resistance to chemotherapeutic agents (Lou et al. 2012; Pasquier et al. 2013; Wang et al. 2018; Desir et al. 2018). Despite the important roles of TNTs in cancer, the regulatory mechanisms and signalling pathways associated with TNT formation are poorly defined, especially in NSCLC.

Studies have mainly focused on the molecular mechanisms involved in stress-mediated TNT formation which occur through the Akt/PI3K/mTOR and p38 MAPK signalling pathways (Zhu et al. 2005; Wang et al. 2011). Actin regulators including the Rho GTPases (CDC42 and Rac1) and M-Sec are known to regulate TNT formation (Gousset et al. 2013; Hase et al. 2009; Schiller et al. 2013). However, the signalling pathways involved in TNT formation are cell line-specific. Minimal studies have investigated the role of exogenous factors and the signalling pathways involved in lung cancer cells despite its presence *in vivo* (Lou et al. 2012).

The hepatocyte growth factor (HGF) signalling axis has been implicated in NSCLC progression (Ichimura et al. 1996; Tretiakova et al. 2011; Olivero et al. 1996). HGF is a pleiotropic cytokine known for exerting cellular effects such as morphogenesis, proliferation and enhanced cellular motility (Ohmichi, Matsumoto, and Nakamura 1996; Singh-Kaw, Zarnegar, and Siegfried 1995; Yanagita et al. 1993; Stoker et al. 1987). Constitutive activation of HGF’s tyrosine kinase receptor (RTK) c-Met, a proto-oncogene (Sonnenberg et al. 1993), induces multiple signalling cascades including the MAPK, PI3K and focal adhesion kinase (FAK) pathways in cancer (Corso, Comoglio, and Giordano 2005; Gherardi et al. 2006; Zhang and Vande Woude 2003). In NSCLC, ligand-dependent c-Met activation occurs through a paracrine route between c-Met expressed cancer cells and stromal HGF or through an autocrine route from cancer cells producing HGF (Masuya et al. 2004; Nakamura et al. 2007).

Crosstalk between β1-integrin with the HGF/c-Met pathways has been widely documented in lung cancer(Barrow-McGee et al. 2016; Ju and Zhou 2013). HGF activation induces integrin clustering and recruits FAK and paxillin which leads to the transduction of downstream MAPK, PI3K and Rho GTPase pathways involved in the biological effects of HGF (Liu et al. 2002; Lai et al. 2000; Ishibe et al. 2004; Ishibe et al. 2003). A major unanswered question centres on whether this signalling axis is involved in TNT formation of A549 cells.

Therefore, the aim of this study was to determine whether HGF/c-Met/β1-integrin signalling axis regulates TNT formation in A549 lung adenocarcinoma cells. This work demonstrates a novel role for the co-activation of HGF/c-Met and β1-integrin in regulating TNT formation in A549 cells *via* paxillin and downstream Arp2/3 complex, MAPK and PI3K pathways.

## Results

### HGF induces the formation of TNT-like structures in A549 lung adenocarcinoma cells

To determine the effect of HGF on TNT formation of A549 cells, white light images were captured on an inverted microscope after 24-hour treatment of HGF (0-700ng/ml)(Fig.1A). HGF induced thin cellular protrusions, narrow at its base and spanning various lengths to connect to distant cells. Therefore, we termed these protrusions TNT-like structures, as they display morphology akin to TNTs. Between 3-30ng/ml, HGF induced A549 cells to scatter and display characteristic elongated morphology. TNT-like structures also began to form as the A549 cells exhibited filopodia-like extensions. At higher concentrations between 100-700ng/ml, TNT-like extensions begin to lengthen across the field of view. To quantify the formation of TNT-like structures, the mean percentage of cells with TNT-like structures, mean number of TNT-like structures per cell and the length of TNT structures were measured. The log concentration-response curve show HGF induces a dose-dependent significant (n=3, \*\*\*\**p*<0.0001) increase in mean percentage, mean number per cell and length of TNT-like structures, with a maximal concentration reached at 100ng/ml plateauing through 300ng/ml and 700ng/ml (Fig.1B-D respectively). However, TNT-like structures were observed at lengths spanning several hundred microns, reaching up to 350μm in our study (Fig.1D). To identify the optimal time point for TNT observation, a 72-hour time course experiment was conducted for HGF at its maximal concentration (100ng/ml) compared to control. The concentration for this time course experiment was determined through additional studies (Supplementary Fig.1A-C). The line graph (Fig.1E-G) displayed a dome-shaped response curve in mean percentage, number per cell and length of TNT-like structures over 72 hours. The maximal timepoint was reached at the 24^th^ hour for two of the parameters: mean percentage (Fig.1E) and mean length (Fig.1G) of TNT-like structures.

**Figure 1:**
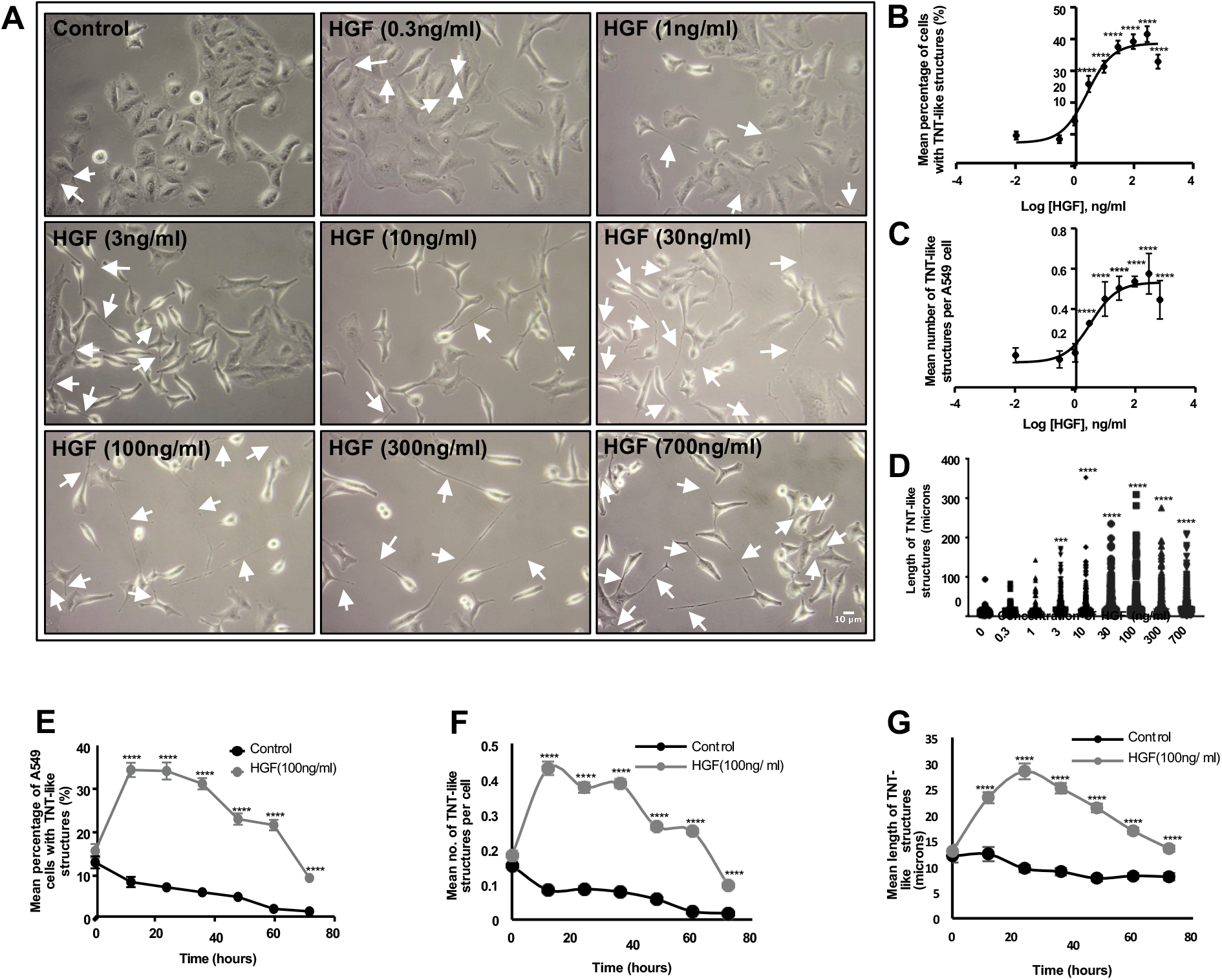
HGF induces TNT-like structures in A549 lung adenocarcinoma cells in a time and concentration dependent manner. (A) Representative white light images demonstrate an increase in TNT-like structures (white arrow) with increasing concentrations of HGF (0-700ng/ml). White light images were captured using 10x objective lens on an inverted microscope. Scale bar: 10μm. The white light images were quantified for TNT-like structures and the HGF log concentration response curve displays the mean percentage (B), number of TNT-like structures per cell (C) and length of TNT-like structures (D). (E-G) The line graphs show the dose-dependent increase in mean percentage (E), number (F) and length (G) of the HGF (100ng/ml)-induced TNT-like structures over a 72-hour time period when compared to control. The maximal effect was observed at the 24th hour timepoint for mean percentage (E) and mean length (G). Values are expressed as mean ±SEM, n=3 with at least 600 cells analysed per condition. \*\*\**p*<0.001 and \*\*\*\**p*<0.0001 when compared to control.

Therefore, the maximal concentration (100ng/ml) and time point (24h) induced by HGF was used for subsequent experiments. We next aimed to determine whether these TNT-like structures were TNTs.

### HGF induces TNT markers F-actin, α-tubulin and M-sec

TNTs structurally comprise of F-actin with microtubules in the thicker regions of TNTs (Rustom et al. 2004; Önfelt et al. 2006). M-sec, has been implicated in the formation of TNTs in multiple cell lines (Hase et al. 2009; Kimura et al. 2016; Barzilai et al. 2016). Confocal imaging and subsequent immunofluorescent labelling with phalloidin, α-tubulin and M-sec confirmed the TNT-like structures contain F-actin throughout with α-tubulin expressed at the thicker regions (Fig.2A) and M-sec (green arrow) localising at the actin-driven protrusion sites of TNTs (Fig.2B). An important morphological characteristic of TNTs includes a lack of adherence to the substratum of the tissue culture surface. Non-adherence of TNTs was assessed through z-stack acquisition; 3D reconstruction and orthogonal view ensured TNTs were non-adherent through passing over another cell or visible only on higher focal planes (Fig.2C). Overall, the TNT-like structures express the characteristic markers of TNTs and are non-adherent which confirms the presence of TNTs in our study.

**Figure 2:**
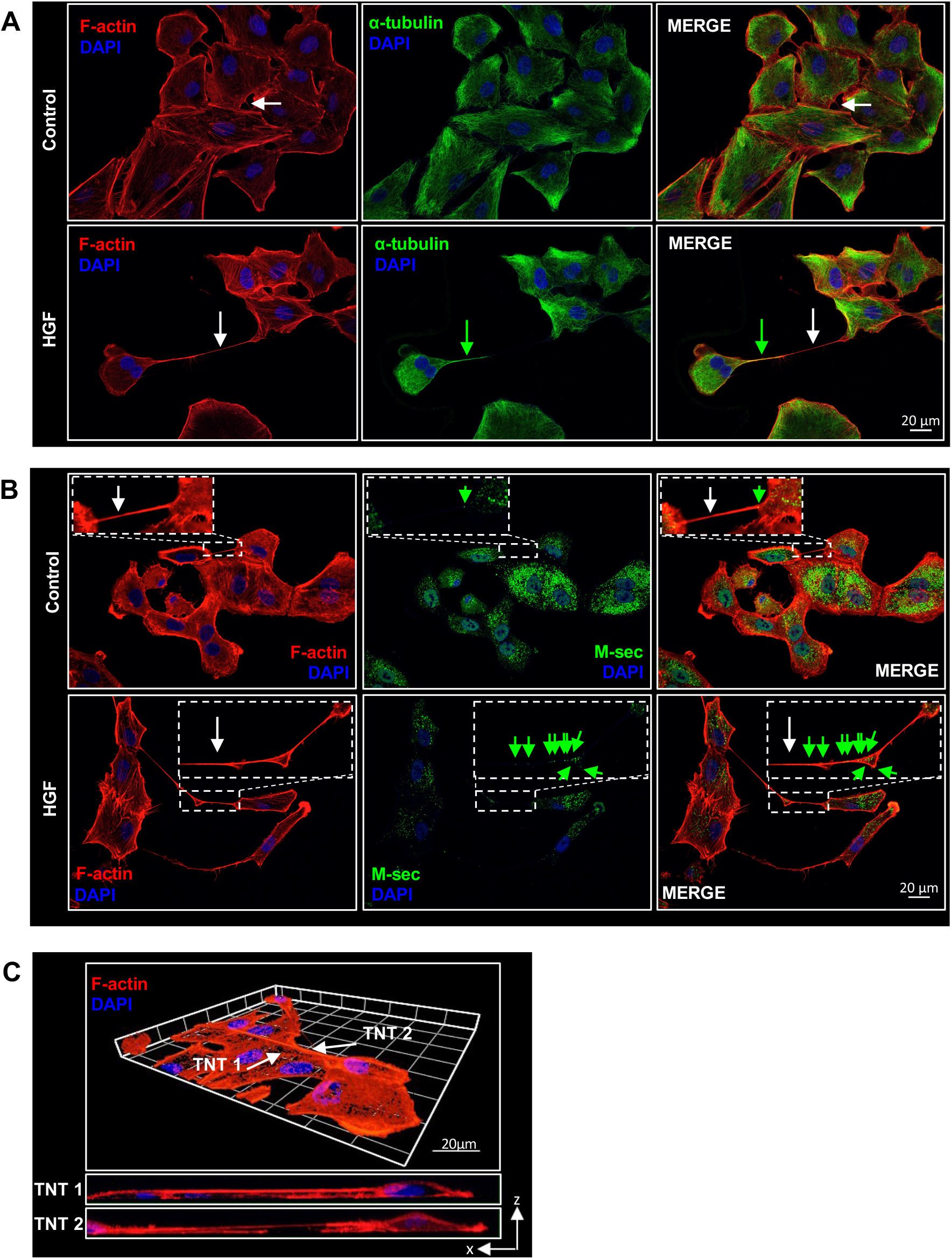
Characterisation of TNT-like structures using immunofluorescent labelling and confocal microscopy. Representative confocal images for control and 24h HGF-treated cells (100ng/ml) labelled with (A) F-actin (red) and α-tubulin (green) and (B) Factin (red) and M-sec (green). M-sec (green arrow) is expressed along F-actin labelled TNTs. White arrows denote TNT structures and the dotted area denotes a zoomed area. (C) Three-dimensional reconstruction Factin positive non-adherent TNT. The threedimensional reconstruction was acquired using the ZEN lite software from 49 sections of Z-stacked images (with Z step of 0.27μm). The maximum xz projection images demonstrates two TNTs hovering above the surface of the substratum. Scale bar, 20μm.

### HGF-induced TNTs transport mitochondria and lipid vesicles

TNTs are known to transport organelles, such as mitochondria, and vesicles (Rustom et al. 2004; Lou et al. 2012). Live-cell confocal microscopy was used to investigate whether HGF-induced TNTs were able to transport mitochondria or vesicles between TNT-connected cells. The time-lapse white light images show two vesicles (white arrow) transferred along a TNT, from a donor cell into the acceptor cell (Fig.3A). To visualise mitochondria transfer, A549 cells were treated with HGF and loaded with a red mitochondrial cytopainter before live-cell confocal imaging was undertaken. The sequence of time-lapse images demonstrates mitochondria transferred along TNT-connected cells (Fig3.B). Furthermore, cells were loaded with either the red mitochondrial cytopainter or DiO lipophilic vesicle dye, before seeding together the two different labelled cell populations and treated with HGF. Fig.3C shows two DiO-labelled vesicles (white arrow) moving along a TNT towards a mitochondria-labelled cell above, furthermore the cell below contains both DiO and mitochondria suggesting prior mitochondria transfer occurred along the TNT. This suggests a unidirectional exchange of organelles between TNT-connected cells.

**Figure 3:**
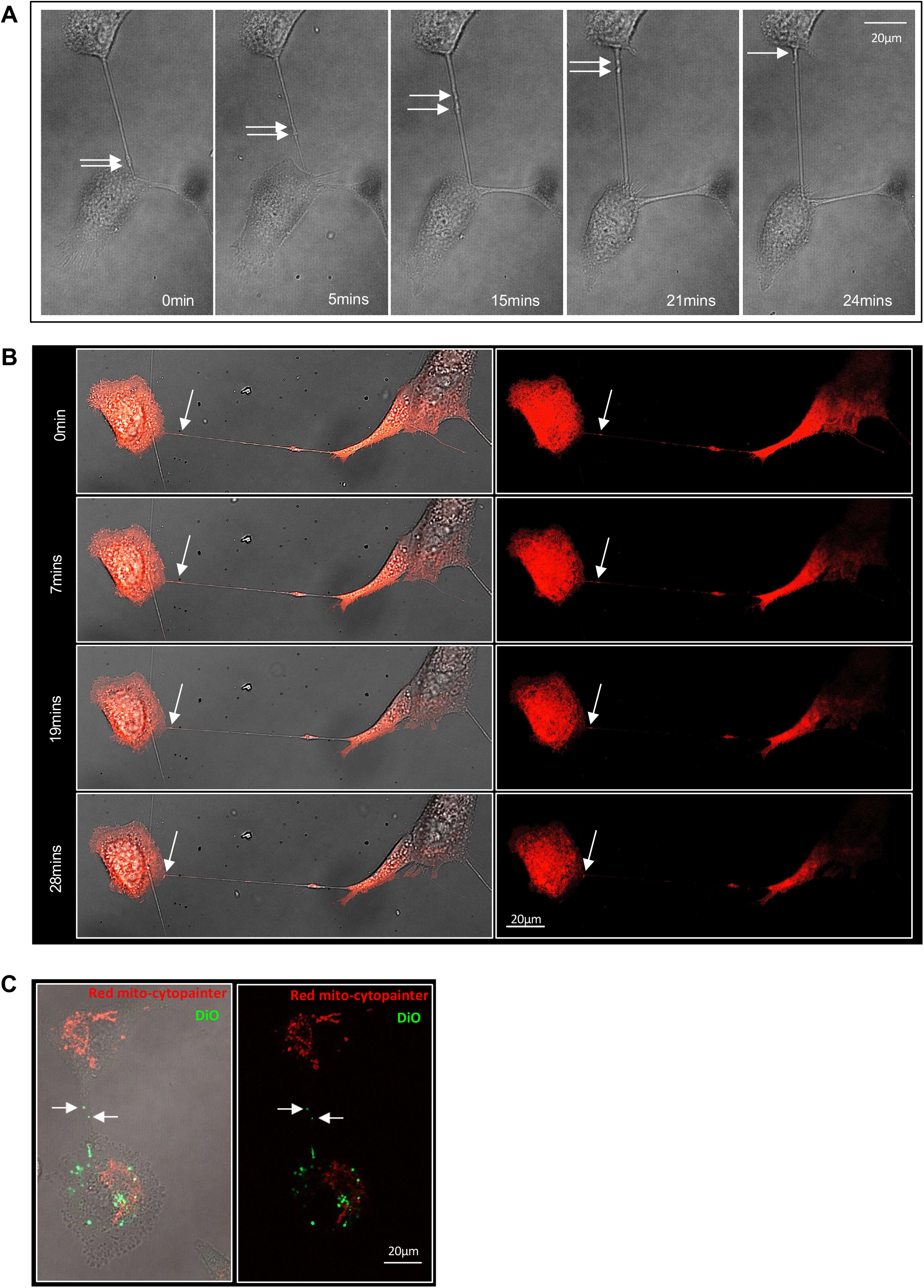
TNTs in A549 cells functionally transmit mitochondria and vesicles. (A) Representative confocal white light images with the indicated time points from a time-lapse movie showing two vesicles (indicated by the white arrows) travelling along a TNT to the cell above. (B) Representative confocal fluorescent images with the indicated time points from a time-lapse movie showing mitochondria transfer along a TNT. A549 cells were loaded with red mitochondrial cytopainter and treated with HGF (100ng/ml) for 24 hours before being imaged on live confocal microscopy. Unidirectional mitochondria transfer along the TNT was observed (indicated by the white arrow). (C) Representative confocal merged light and fluorescence image of two DiO labelled vesicles (indicated by the green arrows) travelling along a TNT towards a red mitochondria-stained cell. Separate populations of A549 cells were loaded with either DiO or red mitochondrial cytopainter and seeded together before HGF treatment and undertaking confocal imaging as above. Unidirectional transfer is suggested through the presence of mitochondria (red) in the DiO (green) labelled cell below. Scale bar, 20μm.

### C-Met receptor regulates TNT formation

To further confirm the role of HGF and its receptor, c-Met, activation modulates TNT formation, a c-Met inhibitor (PHA-665752;0.25μM) was used. The presence of the c-Met inhibitor prevented the effects of HGF, reverting cells to an epithelial cell morphology whilst remaining in cell colonies and fewer TNTs were also observed compared to the HGF DMSO treated cells (shown by white arrows)(Fig.4A). Quantification of the white light images further validated the inhibition of HGF-induced TNT formation with the c-Met inhibitor. A significant (n=3, \*\*\*\**p*<0.0001) decrease was observed in the mean (Fig.4B), mean number per cell (Fig.4C) and length (Fig.4D) of HGF-induced TNTs in the presence of the c-Met inhibitor compared to DMSO. Immunofluorescent labelled TNTs confirmed the presence of TNTs with F-actin (red arrow) expressed throughout the TNT length and M-sec (pink arrow) expressed at the protrusion site of TNTs. Furthermore, the c-Met inhibitor, used at a higher concentration (1μM), significantly decreased the mean percentage (n=3, *\*p*<0.05) and mean number (n=3, *\*\*\*p*<0.001) of control TNTs (Supplementary Fig.2B&C).

**Figure 4:**
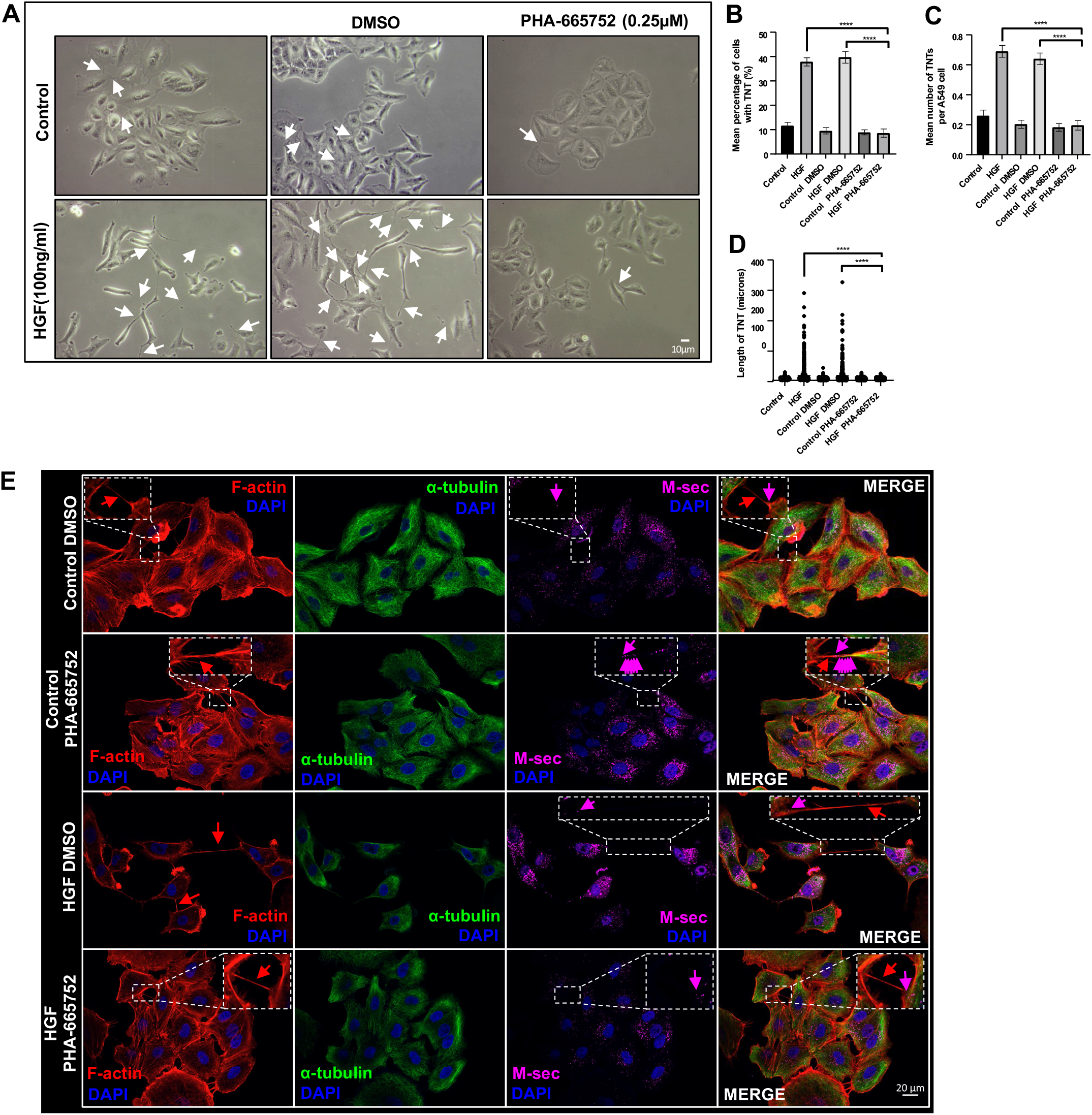
C-Met receptor mediates TNT formation in A549 cells. (A) Representative white light images of control or HGF-treated cells (24 hrs)(100ng/ml) pre-treated with PHA-665752(0.25μM) or DMSO. A decrease in TNTs (white arrow) in both control conditions (DMSO and PHA-665752) was observed. White light images were captured using 10x objective on an inverted microscope. Scale bar, 10μm. (B-D) There was a significant decrease in HGF-induced TNTs in mean (B), number (C) and length (D) of TNTs in the presence of PHA-665752 compared to HGF DMSO treated cells. (E) Representative immunofluorescence confocal images of control or HGF-treated cells (24hrs)(100ng/ml) pre-treated with PHA-665752 (0.25μM) or DMSO and labelled with F-actin (red), α-tubulin (green) and M-sec (pink). There were fewer F-actin-positive HGF-induced TNTs (red arrow) observed in the presence of PHA-665752 compared to DMSO. M-sec labelling (pink arrow) at the protrusion site of TNTs confirmed the presence of TNTs. The dotted region denotes a zoomed area. Scale bar, 20μm. Values are expressed as mean ± SEM, n=3 with at least 900 cells analysed per condition. ****p<0.0001 when comparing HGF + PHA-665752-treated cells with either HGF or HGF + DMSO-treated cells.

### β1-integrin and c-Met receptor crosstalk is involved in TNT formation

Since β1-integrin could be implicated in TNT formation, the next step was to determine whether crosstalk between HGF/c-Met and β1-integrin signalling axis would modulate TNT formation in A549 cells. White light images showed a decrease in TNTs and cells displaying an epithelial morphology in the presence of the functionally blocking β1-integrin antibody compared to the IgG in HGF-stimulated cells (Fig.5A). Further quantification of the white light images demonstrated a significant decrease (n=3, \*\*\*\**p*<0.0001) in mean percentage (Fig.5B), the mean number of TNT per cell (Fig.5C) and length (Fig.5D) in HGF treated cells. These results demonstrate a novel role between β1-integrin and HGF/c-Met in regulating TNT formation in A549 cells.

**Figure 5:**
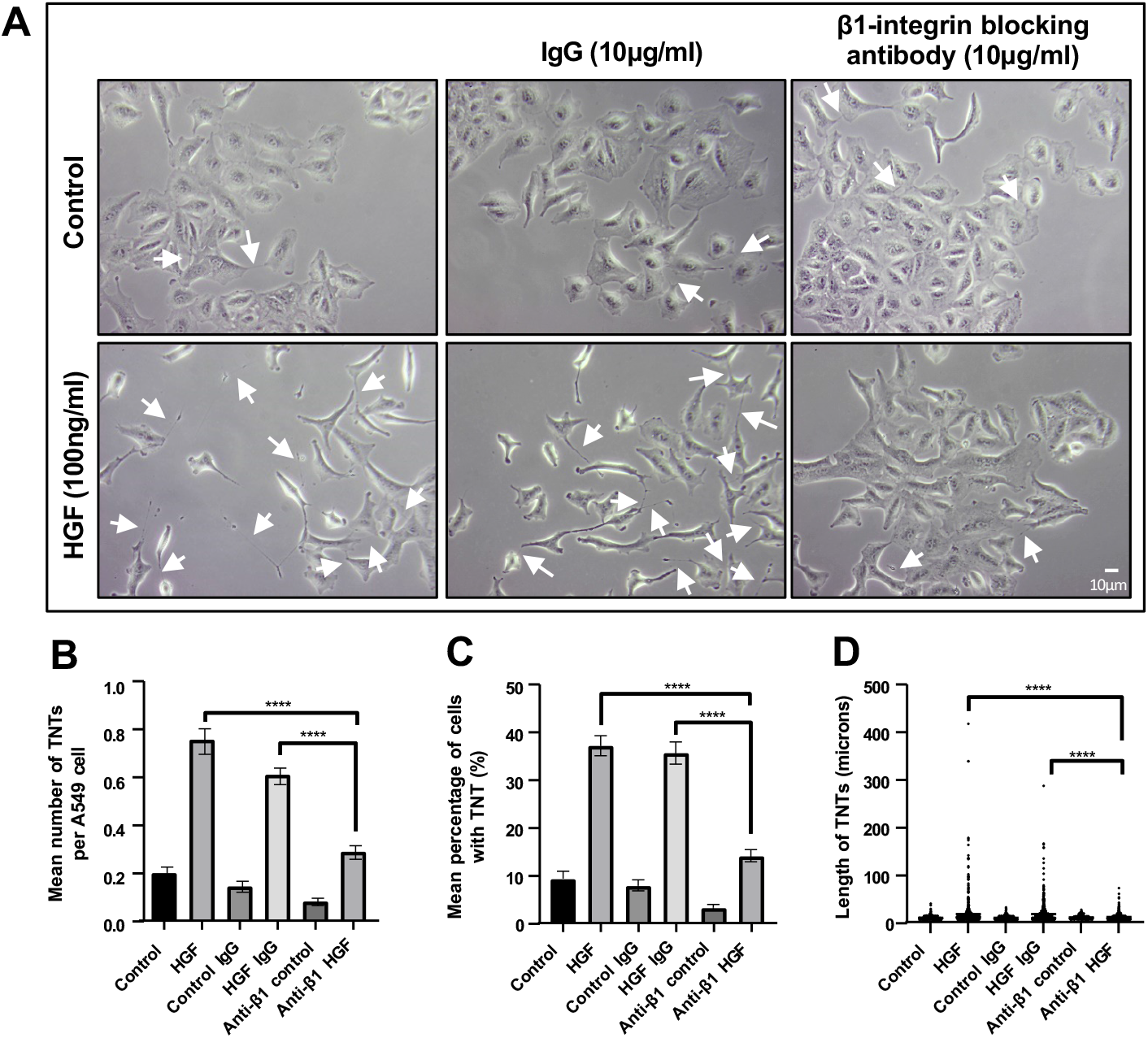
β1-integrin mediates the HGF-induced TNT formation in A549 cells. A549 cells were pre-treated with either IgG or a functionally blocking an anti-β1-integrin blocking antibody (10μg/ml) for 30min before HGF (100ng/ml) treatment for 24hrs. (A) Representative white light images show a decrease in TNTs (white arrow) and colonies of epithelial cell morphology observed in the presence of β1-integrin antibody when A549 cells were treated with HGF compared to its IgG. White light images were captured using 10x objective on an inverted microscope. Scale bar, 10μm. The white light images were quantified for TNTs for the mean percentage (B), mean number (C) and length (D) of TNTs. There was a significant decrease in HGF-induced TNTs quantified in the presence of the β1-integrin antibody compared to HGF DMSO (B-D). Values are expressed as mean ± SEM, n=3 with at least 600 cells analysed per condition. \*\*\*\**p*<0.0001 when comparing between anti-β1-integrin antibody + HGF-treated cells with either HGF or HGF + DMSO-treated cells.

### C-Met, β1-integrin and paxillin are novel components of TNTs

Following our findings demonstrating HGF/c-Met and β1-integrin in regulating TNT formation, the localisation of c-Met and β1-integrin in TNTs were next assessed through immunofluorescent labelling. In untreated cells, c-Met and β1-integrin localised at the cell membrane however HGF treatment induced cytoplasmic and nuclear internalisation of c-Met as previously observed (Ménard, Parker, and Kermorgant 2014). β1-integrin (green arrow) was expressed throughout the entire length of TNTs whilst c-Met (red arrow) was expressed at the thicker portion of the TNT length (Fig.6A). Furthermore, c-Met and β1-integrin co-localised at the protrusion site of TNTs observed in control and HGF-treated cells (yellow arrow)(Fig6.A). To assess the non-adherence of the c-Met and β1-integrin labelled TNTs, z-stack acquisition was obtained to visualise an aerial and xz orthogonal view of the TNT (Fig.6B). The xz orthogonal view shows a c-Met and β1-integrin positive TNT nonadherent to the substratum, which confirms the presence of a TNT.

**Figure 6:**
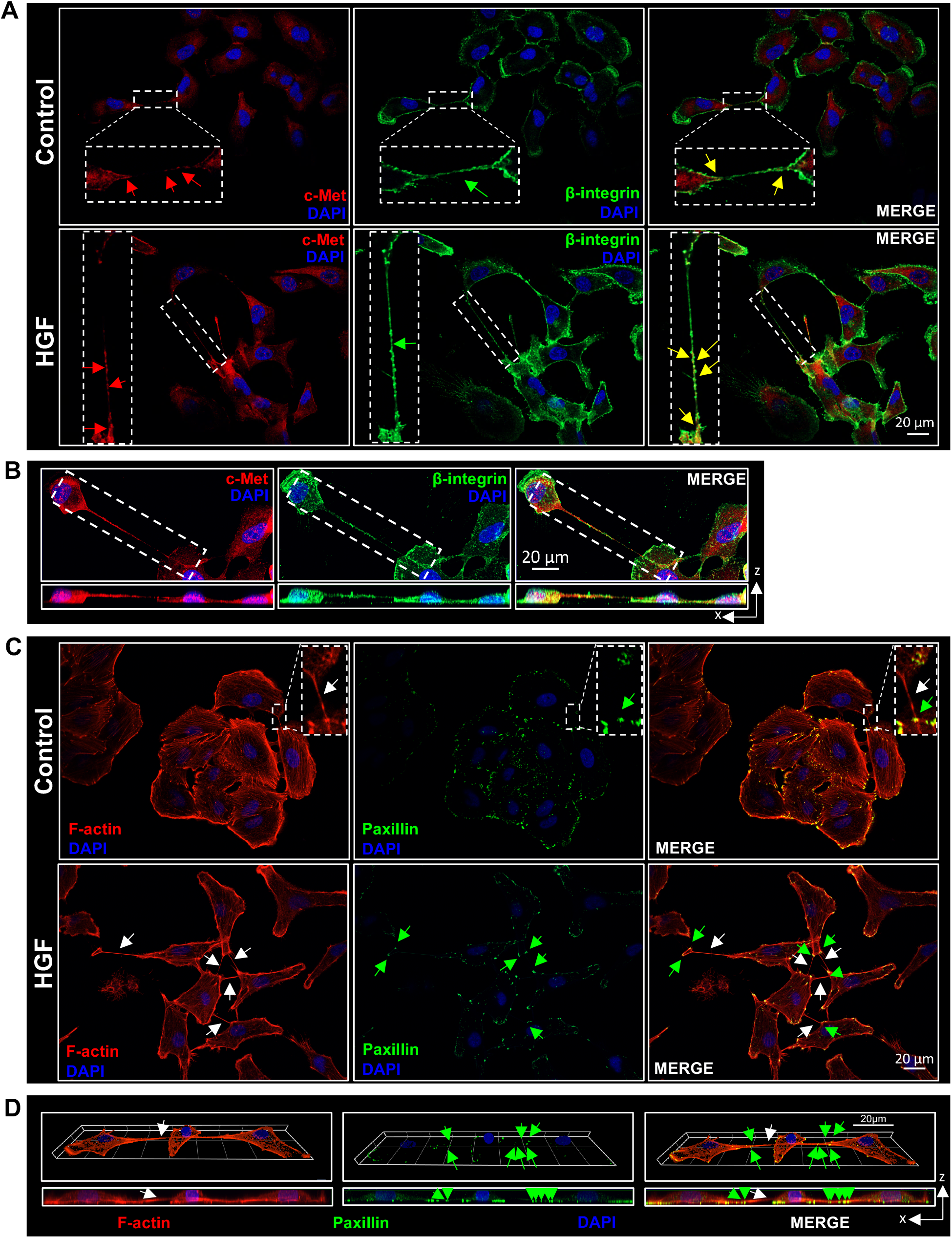
TNTs express novel components c-Met, β1-integrin and paxillin. (A) Representative confocal images for immunofluorescent labelled TNTs with c-Met (red) and β1-integrin (green) in control and HGF-treated A549 cells. C-Met and β1-integrin co-localised along the TNTs (yellow arrow). (B) An example of a non-adherent TNT expressing c-Met (red) and β1-integrin (green). The aerial and orthogonal xz projection image was obtained using ZEN lite software from 39 sections of Z-stacked images (with Z step of 0.27μm). (C) Representative confocal images for immunofluorescent labelled TNTs with paxillin (green) and F-actin (red) in control and HGF-treated A549 cells. Paxillin is expressed at the protrusion site of the control and HGF-induced TNTs (green arrow). The white arrows denote a TNT structure, and the dotted regions denote a zoomed area. (D) An example of a non-adherent F-actin labelled TNT (white arrow) with paxillin (green arrow) expressed at the protrusion site at both ends of the TNT. The 3D reconstruction was acquired using the ZEN lite software from 39 sections of Z-stacked images (with Z step of 0.27μm). The maximum xz projection image below demonstrates the TNT hovering above the surface of the substratum. Scale bar, 20μm.

C-Met and β1-integrin activation has been shown to increase the tyrosine phosphorylation and thus association of paxillin *via* FAK, acting as a point in downstream convergence between HGF/c-Met and β1-integrin signalling (Liu et al. 2002; Crowe and Ohannessian 2004; Schaller et al. 1995; Ishibe et al. 2003). Therefore, paxillin expression was assessed through immunofluorescent labelled TNTs. Paxillin (green arrow) localised at the protrusion site of the F-actin-positive TNTs in control and HGF-treated cells (Fig.6C). Non-adherence was also demonstrated through xz orthogonal view of paxillin expressing TNTs (Fig.6D).

### Arp2/3 complex, MAPK and PI3K pathways mediate TNT formation in A549 cells

To determine the downstream pathways involved in the HGF/c-Met/β1-integrin signalling axis, inhibitors of associated pathways were used. Arp2/3 complex is activated downstream of CDC42/N-WASP and Rac1/WAVE pathways and organises actin filaments into a branched network and is known to be involved in TNT formation (Hanna et al. 2017). The white light images showed a decrease in HGF-induced TNTs (white arrow) in the presence of the Arp2/3 complex inhibitor (CK-666;10μM) when compared to the HGF DMSO treated cells (Fig.7A). The presence of the Arp2/3 complex inhibitor induced a partial yet significant inhibition of the HGF-induced TNTs in mean percentage (Fig.7B)(n=3, \*\*\**p*<0.001), and mean number of TNTs (Fig.7C)(n=3, \*\**p*<0.01) but not in mean length of TNTs when compared to the HGF DMSO treated cells. However, the Rac1 inhibitor (6-Thio-GTP;10μM)(Supplementary Fig.3A-D) and CDC42 inhibitor (ML-141;10μM)(Supplementary Fig.3E-H) did not significantly inhibit the HGF-induced TNT formation when compared to the HGF DMSO treated cells. Furthermore, the ROCK inhibitor (Y27632;5μM) significantly (n=3, \*\*\*\**p*<0.0001) increased the mean (Supplementary Fig.3J), number (Supplementary Fig.3K) and length (Supplementary Fig.3L) of HGF-induced TNTs compared to the HGF DMSO treated cells. The white light images showed the presence of the MAPK (PD98059;20μM)(Fig7.E) and PI3K (LY294002;50μM)(Fig.7I) pathway inhibitors decreased the HGF-induced TNTs (white arrows) and caused the HGF-treated cells to revert to an epithelial cell morphology, similar to the morphology observed in the DMSO control cells. The MAPK and PI3K pathway inhibitors significantly (n=3, \*\*\*\**p*<0.0001) decreased the HGF-induced TNTs in mean percentage (Fig.7F)(Fig.7J) and mean number of TNT per cell (Fig.7G)(Fig.7K) when compared to the HGF DMSO treated cells. The PI3K inhibitor significantly decreased (n=3, *p*<0.001) the mean length of HGF-induced TNTs (Fig.7L) but the MAPK inhibitor had no effect on the HGF-induced TNT length when compared to the HGF DMSO-treated cells (Fig.7H). Moreover, the presence of PD98059 (40μM) alone at a higher concentration, significantly decreased the mean percentage (n=3, \*\**p*<0.01) and mean number (n=3, \*\*\*\**p*<0.0001) of TNTs when compared to control DMSO-treated cells (Supplementary Fig.4B&C). A higher concentration of LY294002 (100μM) also significantly (n=3, \*\*\**p*<0.001) decreased the mean percentage of TNTs when compared to control DMSO-treated cells (Supplementary Fig.4F).

**Figure 7:**
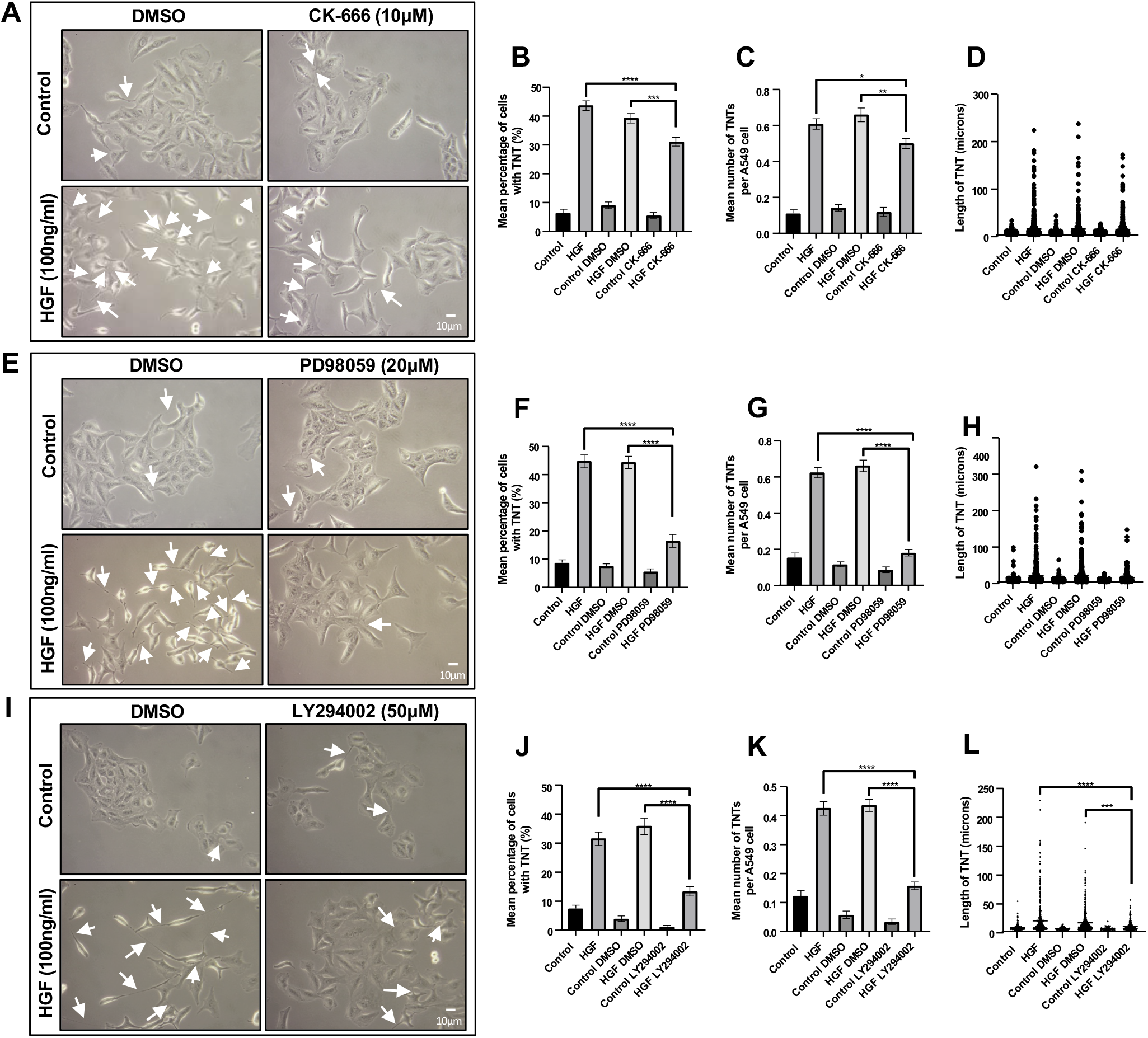
HGF induces TNTs *via* the MAPK, PI3K and Arp2/3 complex pathways. A549 cells were pre-treated with inhibitors associated with the HGF/c-Met downstream pathways or vehicle control for 30mins before HGF treatment (100ng/ml)(24hrs). (A) Representative white light images of control or HGF-treated cells in the presence of CK-666 (10μM) or DMSO. White arrow denotes a TNT structure. White light images were captured using 10x objective on an inverted microscope. Scale bar, 10μm. (B-D) CK-666 significantly decreased the mean (B) and number (C) but not the length (D) of HGF-induced TNTs compared to HGF DMSO treated cells. (E) Representative white light images of control or HGF-treated cells in the presence of PD98059 (20μM) or DMSO. (F-H) PD98059 significantly decreased the mean (F) and number (G) but not the length (H) of HGF-induced TNTs compared to HGF DMSO treated cells. (I) Representative white light images of control or HGF-treated cells in the presence of LY294002 (50μM) or DMSO. (J-L) LY294002 significantly decreased the mean (J), number (K) and length (L) of HGF-induced TNTs compared to HGF DMSO treated cells. Values expressed as mean ± SEM, n=3 with at least 900 cells analysed per condition. \**p*<0.05, \*\**p*<0.01, \*\*\**p*<0.001 and \*\*\*\**p*<0.0001 when comparing between HGF + inhibitor-treated cells with either HGF or HGF + DMSO-treated cells.

### Paxillin regulates the HGF-induced TNT formation

Paxillin is recruited during the co-activation of HGF/c-Met/β1-integrin whilst regulating the downstream signalling pathways. Therefore, to confirm the role of paxillin in TNT formation, A549 cells were transfected with a paxillin siRNA to achieve knockdown. Western blot analysis confirmed significant downregulation of paxillin expression in the A549 cells transfected with the siRNA for paxillin compared to a non-targeting siRNA (Fig.8A). Moreover, white light images revealed a decrease in HGF-induced TNTs when A549 cells were transfected with a siRNA for paxillin whereas the cells transfected with the non-targeting siRNA displayed elongated morphology and an increase in HGF-induced TNTs (Fig8.B). There was a significant decrease in the mean percentage (n=3, \*\*\*\**p*<0.0001)(Fig.8C), the mean number of TNTs (n=3, \*\*\*\**p*<0.0001)(Fig.8D) and mean length (n=3, \**p*<0.05)(Fig.8E), of HGF-induced TNTs when transfected with the paxillin siRNA compared to the cells transfected with the non-targeting siRNA.

**Figure 8:**
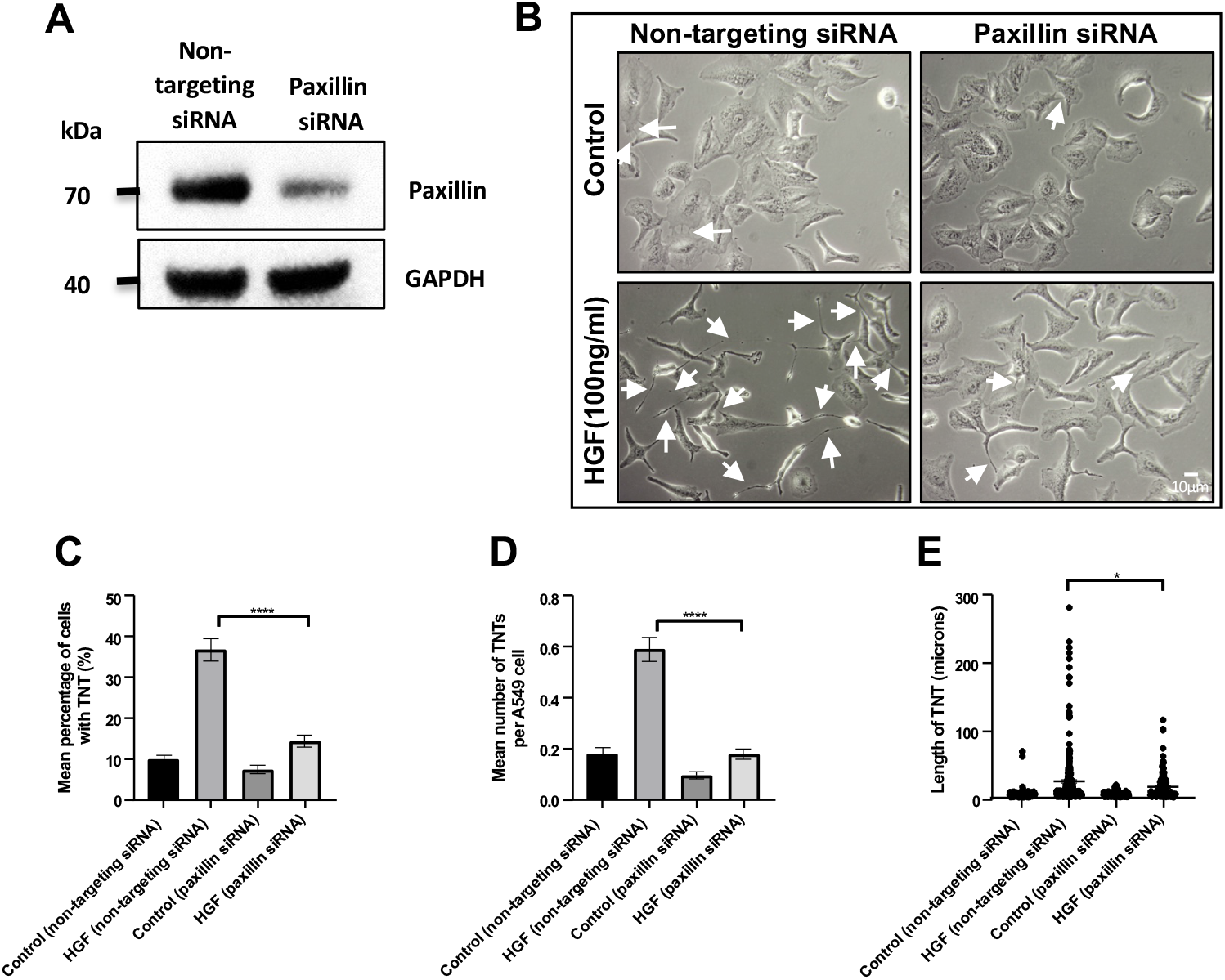
Paxillin regulates HGF-induced TNT formation. (A) Western blot shows knockdown of paxillin in A549 cells after 48h transfection with paxillin siRNA. (B) Representative white light images of A549 cells transfected with paxillin siRNA or non-targeting siRNA for 48h before 24h HGF treatment (100ng/ml). (C-E) There was a significant decrease in mean percentage (C), number (D) and length (E) of TNTs between HGF-treated cells transfected with paxillin siRNA compared to cells transfected with non-targeting siRNA. Values are expressed as mean ± SEM, n=3 with at least 500 cells analysed per condition.\**p*<0.05 and \*\*\*\**p*<0.0001 when comparing between HGF-treated cells transfected with siRNA paxillin and HGF-treated cells transfected with a non-targeting siRNA.

## Discussion

This work highlights a novel role for HGF/c-Met/β1-integrin signalling axis in inducing TNT formation in A549 cells. The TNTs observed demonstrated characteristic non-adherence and expressed F-actin, α-tubulin and M-sec whilst engaging in bidirectional transport of mitochondria and vesicles. Furthermore, c-Met and β1-integrin co-localised at the protrusion site of TNTs and along the TNT length, revealing c-Met and β1-integrin as novel components of TNTs. Crosstalk activation between HGF/c-Met and β1-integrin triggered the downstream PI3K, MAPK and Arp2/3 complex pathways to induce TNT formation. Lastly knockdown of paxillin, through siRNA transfection, inhibited HGF-induced TNTs thus highlighting the central role of paxillin bridging between HGF/c-Met/β1-integrin activation and the downstream signalling pathways involved.

HGF stimulation showed an increase in TNT formation in a time and concentration-dependent manner. TNTs having been observed previously in A549 cells (Wang et al. 2021; Dubois et al. 2018; Kumar et al. 2017; Wang et al. 2012) and in resected human lung adenocarcinoma tissue (Lou et al. 2012). In our study, HGF induced TNTs of up to several hundred microns in length (200-400μm) whereas the maximum TNT length observed in A549 cells was reported at 131μm (Wang et al. 2021). Furthermore, we showed F-actin and α-tubulin as components of TNTs which aligns with previous findings in the cytoskeletal composition of A549 cells (Kumar et al. 2017; Dubois et al. 2018; Wang et al. 2012; Wang et al. 2021). Our study also showed M-sec expression in TNTs which further support the presence of TNTs as M-sec is a known marker of TNT formation in various cell lines (Hase et al. 2009; Kimura et al. 2016). Live-cell imaging also revealed a bidirectional transfer of mitochondria and DiO labelled vesicles along TNTs. Bidirectional transfer of mitochondria and vesicles has also been observed in mesothelioma cells (Lou et al. 2012). The ability to transport large organelles are a unique and determining feature of TNTs. Several studies have alluded to TNT-mediated mitochondria transfer as part of a survival mechanism (Wang and Gerdes 2015; Caicedo et al. 2015; Lin et al. 2019). A549 cells are able to transfer healthy mitochondria to other cells suffering drug-induced mitochondrial loss, thus restoring aerobic metabolism and delaying apoptosis (Spees et al. 2006). Similarly, when PC12 cells were exposed to UV radiation, TNT-mediated mitochondria transfer from healthy cells also rescued cells from undergoing apoptosis (Wang and Gerdes 2015). TNT transfer of mitochondria was also linked to an increase in invasion in bladder cancer cells. Furthermore, when implanted in nude mice, the tumours developed a larger vascularisation network contributing to larger tumours (Lu et al. 2017). Therefore, HGF-induced organelle transfer may be a mechanism to rescue apoptotic cells or alter the phenotypic cellular transformation to achieve invasion and metastasis within the TME.

Furthermore, our study has identified c-Met and β1-integrin as novel components of TNTs, colocalising and engaging in a novel crosstalk activation to regulate TNT formation in A549 cells. Amongst the various mutationally upregulated EMT-inducing-cytokines present in the TME, only EGF and its receptor signalling has been widely studied in TNT formation (Cole, Dahl, and Cowden Dahl 2021; Wang et al. 2011; Hanna et al. 2019). Interestingly, using a higher concentration of the c-Met inhibitor significantly decreased the percentage and number of TNTs in the absence of exogenous HGF treatment. This presents a role for c-Met receptor activation in basal TNT formation. Therefore a paracrine and autocrine route of HGF/c-Met activation, which has been shown previously (Masuya et al. 2004; Nakamura et al. 2007), may be present in regulating NSCLC TNT formation. Other studies have highlighted the importance of the cross-activation between HGF/c-Met and β1-integrin in mediating NSCLC anchorage-independent survival (Barrow-McGee et al. 2016), chemoresistance (Ju and Zhou 2013), metastasis and invasion (Jahangiri et al. 2017). However, our study unravelled a novel non-adhesive role for β1-integrin in the HGF/c-Met-induced TNT formation. Interestingly, a study in murine B lymphocytes showed inhibition of β1-integrin alone had no effect on TNT formation however, inhibition of both α5β1 integrin subunits resulted in the inhibition of TNT formation (Osteikoetxea-Molnár et al. 2016). The difference in these findings highlights the distinct regulatory role of the integrin subunits in regulating TNT formation in different cell types. Moreover, integrins localise at focal adhesion sites and are activated upstream of the signalling pathways that regulate changes in the actin cytoskeletal dynamics (Case and Waterman 2015). FAK has been shown to regulate TNT formation *via* upregulating MMP-2 metalloprotease production in squamous cell carcinoma (Sáenz-de-Santa-María et al. 2017). Breast cancer cells cultured on fibronectin and collagen I also showed an increase in TNT formation (Franchi et al. 2020). Overall, this suggests the dynamic changes occurring in cellular adhesion and ECM remodeling underly the cytoskeletal reorganisation required in TNT formation.

In this study CDC42 and Rac1 inhibition did not inhibit the HGF-induced TNT formation (despite higher concentrations being used-data not shown) this is in contrast to other studies performed in macrophages (Hanna et al. 2017). Despite TNTs being actin-rich structures, there is increasing evidence to suggest that the Rho GTPase pathways play a more complex role in TNT formation dependent on the different cell types. Although CDC42 activation has been proposed in the formation of TNTs in immune cells (Hanna et al. 2017; Arkwright et al. 2010), CDC42 inhibition only minimally decreased the number of TNTs in LST1-induced TNT formation (Schiller et al. 2013). The same study concluded that the M-Sec mediated LST1/RalA/exocyst complex and LST1/RalA/filamin pathway contributed towards the regulation of TNT formation (Schiller et al. 2013). Rac1 has been proposed to sustain existing TNTs instead of playing a role in their formation (Hanna et al. 2017). As the molecular mechanisms underlying TNT formation differ between cell types, our study indicates the Rho GTPases CDC42 and Rac1 are not involved in the HGF-induced TNT formation in A549 cells. We also observed ROCK inhibition promoted an increase in HGF-induced TNTs. Similarly, other studies also showed ROCK inhibition induced an increase in TNT formation *via* Myosin II controlled Factin regulation (Scheiblich et al. 2021; Jana et al. 2022)

MAPK and PI3K inhibition also significantly decreased both basal and HGF-induced TNTs in our study, thus highlighting both signalling pathways as the main regulatory mechanism of TNT formation in A549 cells. Although studies continue to unravel the mechanisms involved in TNT formation, the mechanisms and pathways involved in NSCLC TNT formation remain poorly defined. The MAPK pathway has been implicated in TNT formation in several studies in various cell lines (Cole, Dahl, and Cowden Dahl 2021; Desir et al. 2019; Zhu et al. 2005). Interestingly, in ovarian cancer cells, the PI3K inhibitor did not inhibit TNT formation whereas inhibition of the MAPK pathway decreased the number of TNTs (Cole, Dahl, and Cowden Dahl 2021). However, the PI3K pathway has been shown to be involved in TNT formation in prostate cancer cells and in astrocytes (Kretschmer et al. 2019; Wang et al. 2011). Thus, highlighting again the different molecular mechanisms underlying TNT formation in different cell types. A partial decrease was observed in the presence of the Arp2/3 complex inhibitor. Similarly in macrophages, partial inhibition of TNT formation was also observed (Hanna et al. 2017). Arp2/3 complex is involved in the formation of branched networks of actin, as most TNTs observed in our study are straight in morphology, there would only be a partial inhibition. Furthermore, Arp2/3 complex is mainly activated upstream by CDC42 stimulated N-WASP or Rac1 stimulated WAVE. However, both CDC42 and Rac1 inhibitors did not decrease the HGF-induced TNTs. There may be crosstalk between both the CDC42 and Rac1 pathways in mediating the HGF-induced TNT formation, alternatively, Arp2/3 complex may be activated *via* an ERK-dependent phosphorylation of WAVE (Mendoza et al. 2011). This mechanism of Arp2/3 complex activation has been observed in EGFR activation with an increase in lamellipodia protrusions (Mendoza et al. 2011).

Our study presents a previously undiscovered role for paxillin in HGF/c-Met/β1-integrin-mediated TNT formation. Paxillin acts as a scaffold protein, recruited during the activation of both c-Met and β1-integrin clustering. Paxillin has been shown to localise at the leading edge of membrane protrusions as part of the early stages of focal adhesion formation (Laukaitis et al. 2001). This further supports the notion of paxillin being involved in the initial stages of TNT formation as it is only expressed at the protrusion site of TNTs. Furthermore, paxillin can also bind to various adaptor proteins and multiple downstream signalling effectors to activate the PI3K, MAPK and Arp2/3 complex pathways (Crowe and Ohannessian 2004; Ishibe et al. 2003; Lai et al. 2000). Thus, paxillin acts as the link between the upstream and downstream players involved in the HGF/c-Met/β1-integrin-mediated TNT formation.

Understanding the complex underlying molecular mechanisms involved in TNT formation serves a wider implication in NSCLC therapy. Unravelling the novel signalling axis between HGF, c-Met and β1-integrin in TNT formation highlights the need for a more personalised targeted approach in NSCLC treatment. Future *in vivo* studies would be needed to characterise the occurrence of TNTs and c-Met expression in lung adenocarcinoma tissue. This would serve as the next step in targeting TNTs in NSCLC.

**Schematic 1:**
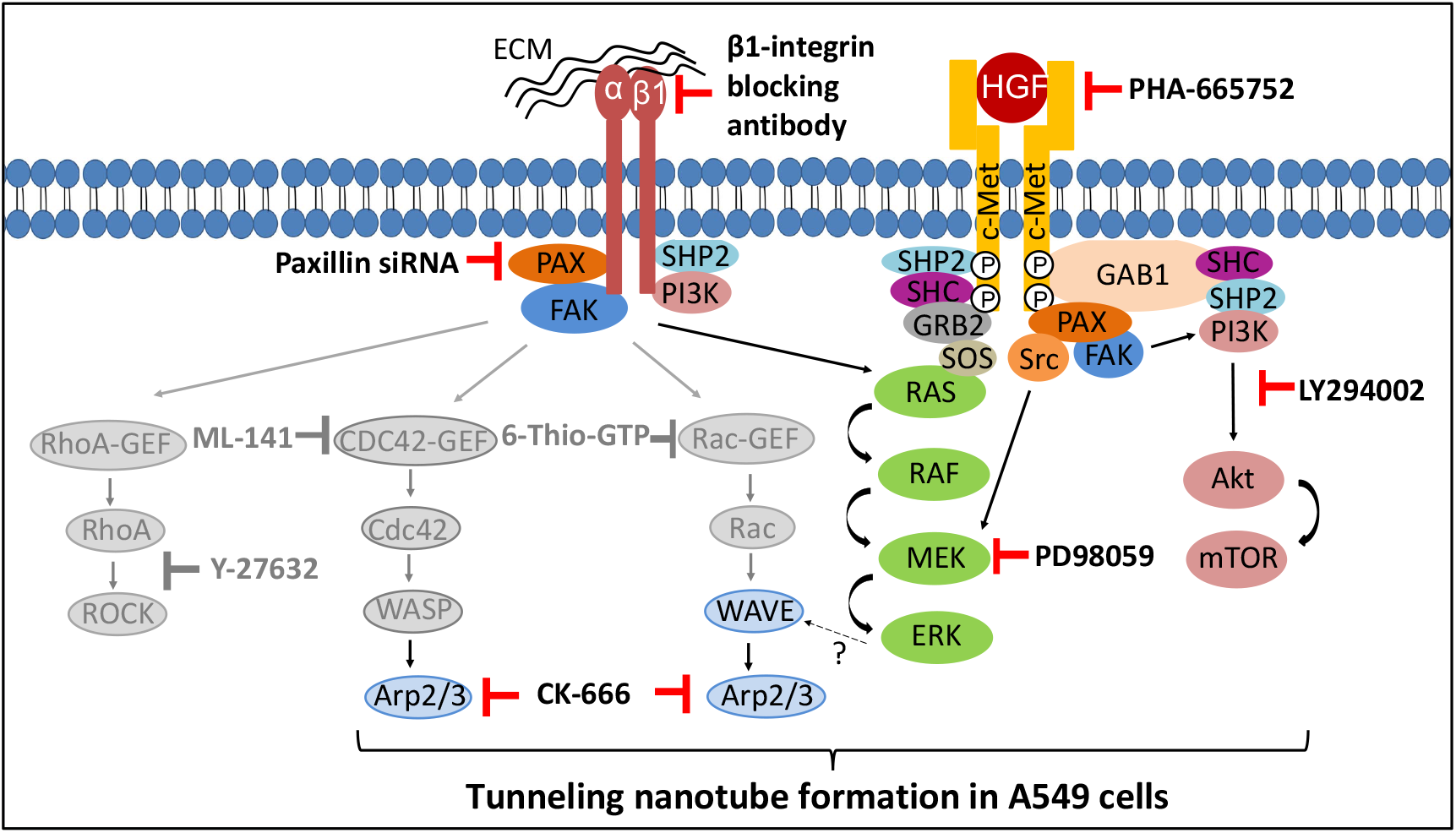
HGF/c-Met and β1-integrin signalling axis drives TNT formation in A549 cells. Paxillin is recruited after HGF/c-Met and β1-integrin activation and acts as a scaffold protein to activate Arp2/3 complex, PI3K and MAPK downstream pathways. The Rho GTPase-associated pathways (Rho, CDC42 and Rac1)(highlighted in grey) did not inhibit the HGF-induced TNT formation. However, the Arp2/3 complex was found to mediate the branched morphology of TNTs which may occur *via* ERK-mediated phosphorylation of WAVE.

## Materials and Methods

### Reagents

*Cytokine and inhibitors;* Purified recombinant human HGF was obtained from Peprotech. The c-Met receptor inhibitor (PHA-665752≤1μM), MAPK pathway inhibitor (PD98059≤40μM), Arp 2/3 complex inhibitor (CK-666≤200μM) and CDC42 inhibitor (ML-141≤20μM) were obtained from Cayman Chemical. The PI3K pathway inhibitor (LY294002≤100μM), Rac1 inhibitor (6-Thio-GTP≤50μM) and ROCK pathway inhibitor (Y27632≤7.5μM) were obtained from Selleck chemicals, Jena Bioscience and Merck respectively. The functionally blocking β1-integrin antibody (≤10μg/ml) and its mouse IgG1 isotype were purchased from Abcam.

*Immunofluorescence;* primary and secondary antibodies; rabbit anti-Met, rabbit anti-TNFAIP2(M-sec), rat anti-α-tubulin was obtained from Invitrogen. Mouse anti-paxillin was obtained from BD Bioscience. Mouse anti-β1-integrin was obtained from Abcam. Alexafluor-conjugated secondary antibodies (488, 568 and 647nm), Rhodamine phalloidin, DiO’; DiOC_18_(3) (3,3’-Dioctadecyloxacarbocyanine Perchlorate) were all purchased from Invitrogen. The mitochondrial staining kit-red fluorescence-cytopainter was purchased from Abcam. Vectashield mounting medium was obtained from Vector Laboratories Ltd and DAPI was obtained from Invitrogen.

### Cell culture

The A549 lung adenocarcinoma cell line was obtained from the American Type Culture Collection (ATCC). The cells were cultured in Dulbecco’s Modified Eagle’s Medium (DMEM)(Gibco) and supplementation of the media required 10% Foetal Bovine Serum (FBS), 200mM L-glutamine, 10 000 U/ml penicillin and 10 000μg/ml streptomycin (Gibco). The cell lines were cultured and maintained at 37°C with CO2 level of 5% in a humidified atmosphere.

A549 cells were seeded at a density of 5×10^3^ cells/well in a 12-well plate in 10% FBS supplemented DMEM media. The cells were left to grow into colonies for 72 hours in a humidified atmosphere previously described before treatment with HGF in 2% FBS supplemented DMEM media for 24h. For inhibitor studies, A549 cells were pre-treated for 30 minutes with the inhibitors of pathways detailed above with their respective vehicle controls. Following pre-treatment, HGF (100ng/ml) in 2% FBS supplemented media was added for 24h before undertaking white light imaging.

### siRNA transfection

The non-targeting siRNA (Qiagen) and paxillin siRNA (Qiagen) were transfected with HiPerfect (Qiagen) as previously described (Porter et al. 2020). 20μM of siRNA/well was transfected on a 6-well plate. Following 48hrs of transfection, cells were trypsinised and a fraction of the cells were used to seed on a 12-well plate for HGF treatment previously described. The remainder of cells were used for Western blot processing to confirm paxillin knockdown. The following paxillin siRNA target sequence was used: ATCCAAAGGCAGAGAACCAAA.

### Immunofluorescence

#### Confocal immunofluorescence microscopy

A549 cells were seeded at a density of 7.5×10^3^cells on No. 0 coverslips (Fisher Scientific) and treated with HGF and its respective pathway inhibitors as previously described. Cells were then fixed with 4% PFA for 10 mins, permeabilised with 0.5% Triton-X for 3.5 min and blocked with 3% BSA before overnight incubation at 4°C with primary antibodies (1:100) in 3% BSA. For immunolabelling of c-Met, cells were fixed with 100% methanol for 10 min, permeabilised with 0.1% Triton-X for 10 mins and blocked with 3% BSA before overnight incubation at 4°C with anti-c-Met primary antibody

(1:100) in 3% BSA. Visualisation of immunolabelled proteins required the respective combination of species-specific Alexafluor-conjugated secondary antibodies (488, 568 and 647nm)(1:200), raised in donkey or goat, co-incubated with rhodamine phalloidin. Immunolabelling protocol for paxillin was followed as per Sastry et al (Sastry et al. 1999). Cells were then imaged with Zeiss LSM980-Airyscan confocal microscope using 40x 1.4 NA oil objective.

#### Live-cell trafficking of mitochondria and vesicles

To visualise vesicle and mitochondria transfer along the TNTs, HGF-pre-treated cells on coverslips were pre-loaded with red mitochondrial cytopainter for 30mins following the manufacturer’s instruction and then washed three times with PBS. Separate cell populations were also preloaded with either DiO (5μM) or red mitochondrial cytopainter for 30mins. Both cell populations were then seeded together at a 1:1 ratio and treated with HGF for 24hrs. Coverslips were mounted on a ludin chamber and enclosed in a humidified atmosphere at 37°C with 5% CO_2_. Real-time acquisition of images was captured at 30 sec-2mins intervals for a duration of 20min-3hrs. Live cells were imaged with Zeiss LSM980-Airyscan confocal microscope using 40x 1.3 NA oil objective.

### TNT image analysis

#### White light image analysis

White light images were acquired with a Zeiss Primovert inverted microscope and GXCAM3EY-5 camera at x10 magnification. At least 10 fields of view per condition were captured with at least 400 cells analysed per condition. Using ImageJ software, the total number of cells were counted and the number of cells with TNTs were counted to obtain the mean percentage of A549 cells with TNT for each field of view. The images were also analysed to obtain the counts for the number of TNT per cell. Individual TNT lengths were also measured in μm, from the narrow initiation point to the end of the visible TNT length.

#### Confocal image analysis

Confirmation of lack of adherence was observed through z-stack acquisitions of F-actin labelled TNTs visible on higher focal planes. Z stack images were acquired at 0.27-1.2μm intervals. The Zenlite software was used to obtain the 3D reconstruction and orthogonal view.

### Western blot

Protein extraction was performed with 4x Laemmli’s Buffer, which comprised of 0.5 M Tris pH 6.8 (12.5%), Glycerol (10%), SDS (2%), Bromophenol Blue (0.08%) and 2-mercaptoethanol (5%) in water. Protein lysates were denatured at 95°C for 5mins before being loaded into a TruPAGE 4-20% precast gradient gel (Sigma) and undergoing SDS-PAGE at 120 V for 2 hours. Separated proteins were subsequently transferred onto PVDF membrane at 30 V for 3 hours. Membranes were blocked with 5% milk/TBST at room temperature for 1 hr before being incubated overnight at 4°C with paxillin (Abcam) and GAPDH (Cell Signaling Technology) primary antibodies diluted in 5% BSA/TBST (1:500 and 1:4000 respectively). After three washes with TBST, membranes were incubated at room temperature for 2 hours in anti-rabbit (1:2000)(Sigma) and anti-mouse (1:4000)(Sigma) secondary antibodies diluted in 5% milk/TBST (1:2000 and 1:4000 respectively). Following incubation, the signals were detected by ECL detection reagents (Amersham).

### Statistical analysis

Experiments were conducted at least three times with at least 500 cells analysed per condition. The data was expressed as the mean ± SEM for log dose-response curves, line graphs, scatter plots and histograms comparing control with inhibitor-treated/siRNA knockdown groups. Determination of significance was assessed appropriately with either one-way ANOVA and Tukey’s multiple comparisons or through unpaired t-test using Graphpad Prism 8. All statistical analysis was significant when *p*<0.05.

## Acknowledgements

The authors thank Prof. Mette Mogensen from the School of Biological Sciences at the University of East Anglia for providing the paxillin and α-tubulin primary antibodies.

The authors gratefully acknowledge the funding from the School of Pharmacy, University of East Anglia and Nurhadibrata Jap.

