## Supplementary Data for "Hepatocyte Growth Factor and β1-integrin signalling axis drives tunneling nanotube formation in A549 lung adenocarcinoma cells"

### Supplementary Information

**A**

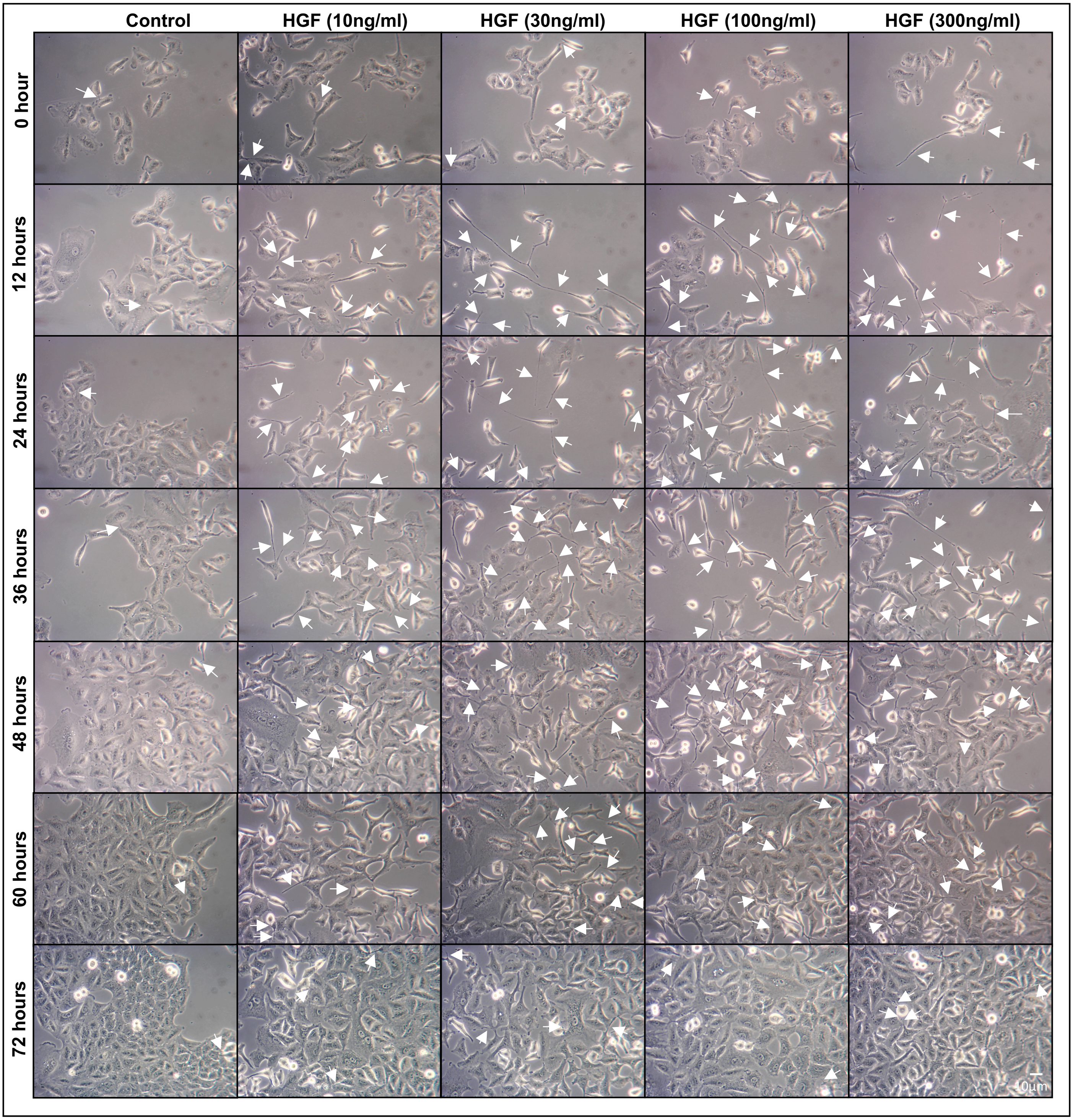

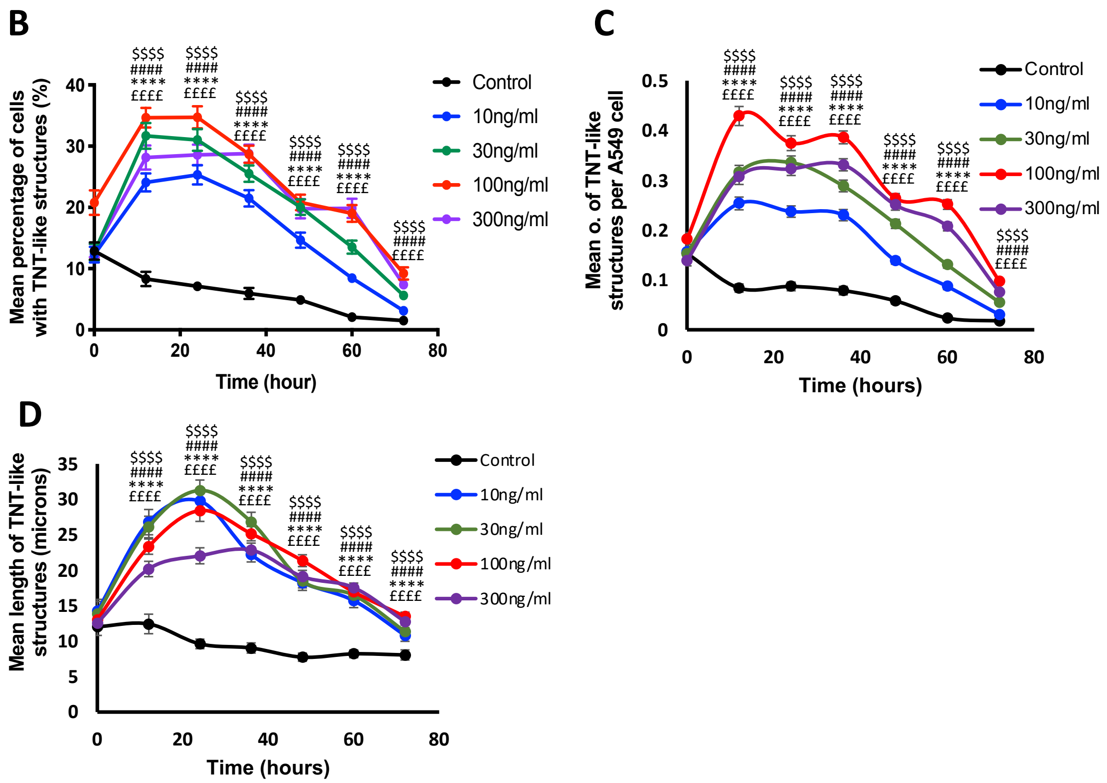


**Supplementary Figure 1 (related to Fig.1):** HGF induces TNT-like structures in A549 lung adenocarcinoma cells in a time dependent manner. (A) Representative white light images demonstrate an increase in TNT-like structures (white arrow) with increasing concentrations of HGF (0-300ng/ml) over 72 hours. White light images were captured using 10x objective lens on an inverted microscope. Scale bar: 10µm. The white light images were quantified for TNT-like structures and the line graphs showed a dose-dependent increase in the mean (B), number (C) and length (D) of the HGF-induced TNT-like structures between control and the different concentrations of HGF-treated cells over a 72-hour time period. The maximal effect was observed at the 24th hour timepoint for the mean percentage and length of the HGF-induced TNT-like structures. Values are expressed as mean ± SEM, n=3 with at least 1500 cells analysed per condition. ££££*p*<0.0001 when comparing control to 10ng/ml*,* ****$p$<0.0001 when comparing control to 30ng/ml, ####*p*<0.0001 when comparing control to 100ng/ml*,* $$$$*p*<0.0001 when comparing control to 300ng/ml across all the timepoints, by one-way ANOVA and Tukey's multiple comparison test.

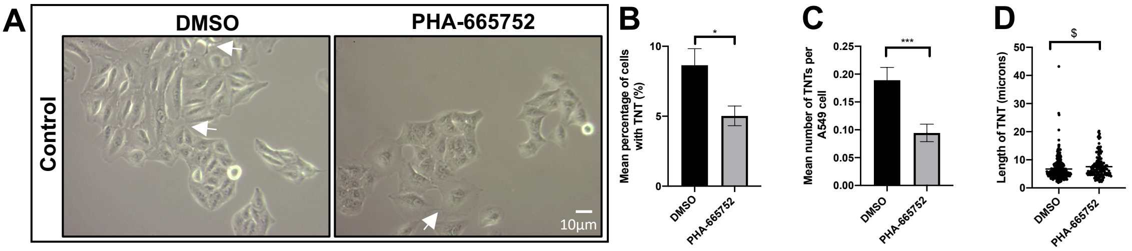


**Supplementary Figure 2 (related to Fig.4):** (A) Representative white light images of A549 cells treated with PHA-665752(1µM) or DMSO. White arrow denotes a TNT structure. White light images were captured using 10x objective on an inverted microscope. Scale bar, 10µm. (B,C) There was a significant decrease in the mean percentage (B) and mean number (C) of TNTs. (D) A significant increase in TNT length was observed in the PHA-6665752 treated cells compared to DMSO. Values are expressed as mean ± SEM, n=3 with at least 400 cells analysed per condition. *$p$<0.05 and ***$p$<0.001 when compared to respective DMSO controls. $$p$<0.05 when PHA-665752 TNT length was compared with DMSO control.

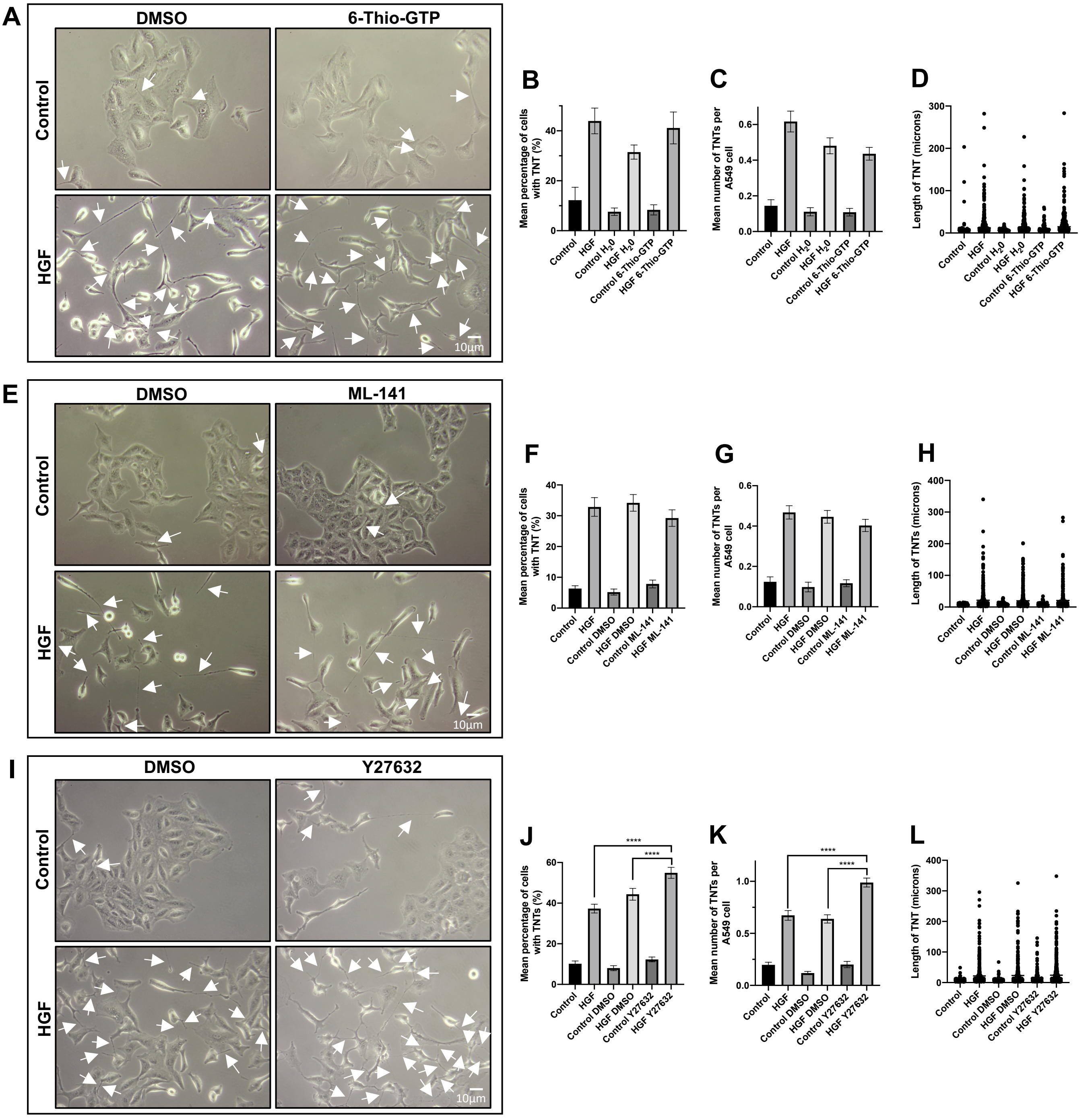


**Supplementary Figure 3: HGF does not induce TNTs *via* the ROCK, Rac1 and CDC42 pathways.** A549 cells were pre-treated with inhibitors associated with the HGF/c-Met downstream pathways or vehicle control for 30mins before HGF treatment(100ng/ml)(24hrs). (A) Representative white light images of control or HGF-treated cells in the presence of 6-Thio-GTP (10µM) or DMSO. (B-D) In the presence of 6-Thio-GTP, there was no significant difference in mean percentage (B), number (C) and length (D) of TNTs compared to HGF $H_20$ treated cells. White arrow denotes a TNT structure. White light images were captured using 10x objective on an inverted microscope. Scale bar, 10µm. (E) Representative white light images of control or HGF-treated cells in the presence of ML-141 (10µM) or DMSO. (F-H) In the presence of ML-141, there was no significant difference in mean percentage (F), number (G) and length (H) of TNTs compared to HGF DMSO treated cells. (I) Representative white light images of control or HGF-treated cells in the presence of Y27632 (5µM) or DMSO. White arrow denotes a TNT structure. (J-L) The presence of Y27632 induced a significant increase in mean percentage (J), number (K) and length (L) of HGF-induced TNTs compared to DMSO HGF-treated cells. Values are expressed as mean ± SEM, n=3 with at least 500 cells analysed per condition. ****$p<0.0001$ when comparing HGF Y27632 treated cells with either HGF treatment or HGF DMSO-treated cells.

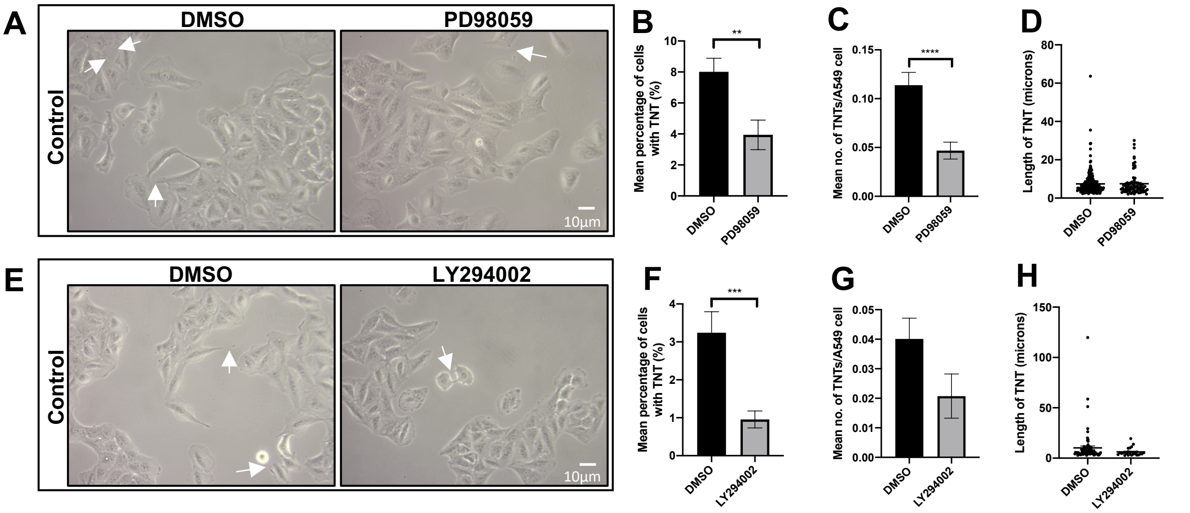


**Supplementary Figure 4 (related to Fig7):** (A) Representative white light images of A549 cells treated with PD98059 (40µM) or DMSO. White arrow denotes a TNT structure. White light images were captured using 10x objective on an inverted microscope. Scale bar, 10µm. (B-D) There was a significant decrease in the mean percentage (B) and mean number (C) but not in length (D) of TNTs. (E) Representative white light images of A549 cells treated with LY294002 (100µM) or DMSO. (F-H) There was a significant decrease in the mean percentage (F) but not in mean number (G) and length (H) of TNTs. Values are expressed as mean ± SEM, n=3 with at least 400 cells analysed per condition. **$p<0.01$, ***$p<0.001$ and ****$p<0.0001$ when compared to respective DMSO controls.
